# A Synthetic Genetic Reversible Feynman Gate in a Single *E.coli* Cell and its Application in Bacterial to Mammalian Cell Information Transfer

**DOI:** 10.1101/2021.08.16.456494

**Authors:** Rajkamal Srivastava, Kathakali Sarkar, Deepro Bonnerjee, Sangram Bagh

## Abstract

Reversible computing is a nonconventional form of computing where the inputs and outputs are mapped in a unique one-to-one fashion. Reversible logic gates in single living cells have not been demonstrated. Here, we created a synthetic genetic reversible Feynman gate in a single *E.coli* cell. The inputs were extracellular chemicals, IPTG and aTc and the outputs were two fluorescence proteins EGFP and E2-Crimson. We developed a simple mathematical model and simulation to capture the essential features of the genetic Feynman gate and experimentally demonstrated that the behavior of the circuit was ultrasensitive and predictive. We showed an application by creating an intercellular Feynman gate, where input information from bacteria was computed and transferred to HeLa cells through shRNAs delivery and the output signals were observed as silencing of native AKT1 and CTNNB1 genes in HeLa cells. Given that one-to-one input-output mapping, such reversible genetic systems might have applications in diagnostics and sensing, where compositions of multiple input chemicals could be estimated from the outputs.

## Introduction

The adaptation of logic circuit principles for building synthetic genetic logic circuits allowed performing several computational operations in living cells^**1–4**^. Some of such examples include implementation of various synthetic genetic logic gates and integrated logic circuits^**1–4**^, counters^**5**^, switches^**6**^, oscillators^**7**^, half and full adders^**8,9**^, multiplexer and demultiplexer^**10**^, and analog-to-digital converters^**11**^. The logical operations used in synthetic biology are based on the conventional logic gates. Such logic gates like AND, OR, XOR, or NOR work in such a way that it irretrievably loses some information during computation^**12**^. Thus, those conventional operations are irreversible in nature.

The reversible computing is a form of an unconventional computing, where the computation is thermodynamically and logically reversible^**12**^. Such computation should be performed in a thermodynamically reversible way, such that the energy associated with a computation operation would not dissipated as heat energy but be reused for the next computing operation. On the other hand, logical reversibility allows reconstructing the previous state of a computing operation from the current computing state. In general, reversible gates consume their own inputs. That means reversible gates do not have higher numbers of inputs than its output. The logical reversibility allowed a one-to-one mapping between its input and output signals and as a result input signals can be identified and traced from its unique output signals^**12**^. Thus, a reversible logic gate does not wipe out any information and a reversible circuit made of reversible gates does not have fanouts and loops resulting equal numbers of input and output wires^**13**^. Reversible logic gates are the heart of quantum computing. Reversible logic gates were mostly implemented using CMOS and photonics^**14,15**^. Further, logical reversibility has been demonstrated with in-vitro biochemical reactions, where the logical reversibility but not thermodynamic reversibility of Fredkin, Toffoli, Feynman and Peres gates have been implemented using in-vitro DNA computation^**16**^ and enzyme catalyzed reactions^**17**^.

A living cell and its cellular processes stay out of equilibrium. Therefore, attaining thermodynamic reversibility in living cells is not possible. However, it may be possible to attain logical reversibility in living cells. Recently we have demonstrated logical reversibility in living cells by implementing Feynman and Fredkin gates in bacteria by using an artificial neural network (ANN) type architecture made from a mixed population of engineered bacteria connected by chemical wires^**18**^. Thus, logical reversibility in living cells was achieved through distributive computing where various populations of cells carried out various component functions and as a mixed population they showed the complete function. However, to expand the capability of reversible logic circuits in living cells, it is important to create reversible logic gates in single cells.

In this work we demonstrated a synthetic genetic reversible Feynman gate in a single *E.coli* cell, where the inputs are small molecule extracellular chemicals IPTG and aTc and the outputs were two fluorescence proteins EGFP and E2-Crimson. We developed a simple mathematical model and simulation to capture the essential features of the genetic Feynman gate and experimentally demonstrated that the circuit was ultrasensitive or ‘digital like’ and the behavior of the circuit matched with the computational simulation. Further, we showed a simple application by creating a bacterial-mammalian intercellular genetic Feynman gate. We demonstrated shRNA-mediated gene silencing in HeLa cells using genetic Feynman gate, where the input information from the bacteria is reversibly transferred to mammalian cells. Here the input signals were applied in the engineered invasive *E.coli* cells and the information were transferred to HeLa cells through shRNAs and the outputs were manifested by repression of two native genes, AKT1 and CTNNB1 in HeLa cells.

## Results and Discussions

### Genetic circuit designs for the Feynman gate

The Feynman gate or controlled NOT gate^**12**^ has two inputs A and B and two outputs P and Q (Fig. 1A). It is clear from its truth table that there is a one-to-one mapping between input and outputs, where the output signals are determined as P = A and Q = A ⊕ B (Fig. 1A). The first input A is a control input line, whereas the other input B is target input line. The target line shows negation operation and can only be performed when control line is set. Otherwise, target line shows no operation. Due to this controlled negation operation Feynman gate is also known as Controlled NOT Gate. We designed a synthetic genetic Feynman gate in a single cell (Fig. 1B). The design consisted of four synthetic promoters. The promoter P_LTATA_ consisted of two lac-operating sites and two tet operating sites. LacR and TetR proteins bind with the lac and tet operating sites respectively^**18**^. Presence of any of the repressor proteins in the cell blocks the binding and movement of RNA polymerase on the promoter and prevents the transcription process from the P_LTATA_. IPTG and aTc, two small molecule inducers, when applied from outside, diffuse in the cell and bind with the LacR and TetR respectively. Those molecular bindings change the conformation of LacI and TetR in such a way that they could not bind with their operating sites any more^**19**^. Thus, the simultaneous presence of the IPTG and aTc activates the transcription. TetR and LacI proteins were constitutively expressed from the engineered chromosome of *E.coli* strain DH5αZ1^**19**^. A mutated version of CI protein^**20**^ (mCI) was placed under this P_LTATA_ promoter. This mCI protein, when expressed could bind with two synthetic promoters P_LAACCC_^**18**^ and P_LCCTT_^**18**^ to prevent the transcription from these two promoters.

**Figure 1.**
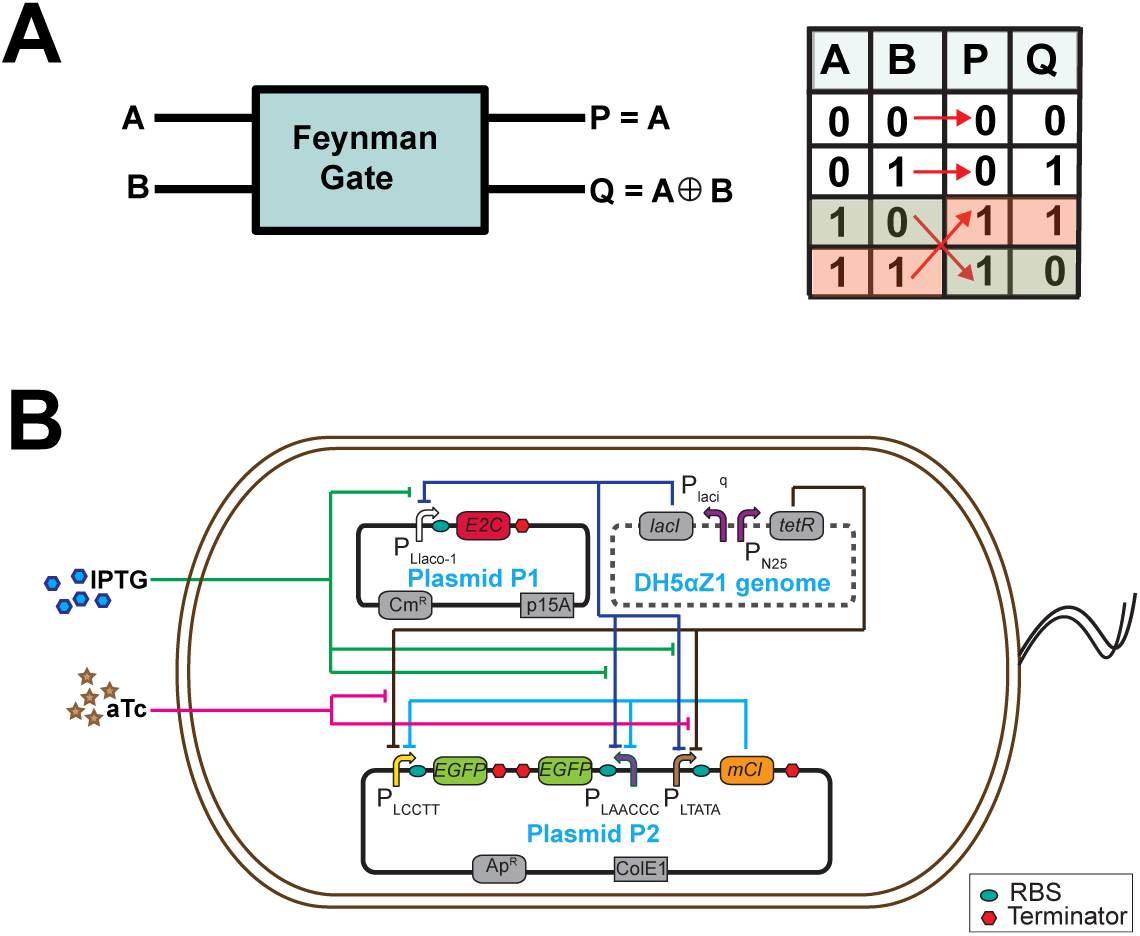
A biological 2×2 Feynman Gate Device. **A)** Block diagram (left) and truth table (right) of electronic Feynman gate associated with two inputs (A and B) and two outputs (P and Q). Red arrows represent input to output mapping. **B)** Synthetic genetic design of Feynman gate in engineered *E. coli* processing two environmental input chemicals (IPTG and aTc)

The promoter P_LAACCC_ consisted of two lac-operating sites and two CI binding sites such that the presence of any of the repressors prevents the transcription process. Similarly, promoter P_LCCTT_ consisted of two tet operating sites and two CI binding and its transcription is inhibited by tetR and mCI. One copy of EGFP gene was placed under each P_LAACCC_ and P_LCCTT_ promoter. We put an E2Crimson gene under an IPTG activated PLlaco-1 promoter. Thus by tuning the extracellular aTc and IPTG, the expression of EGFP and E2-crimson can be logically controlled. The bioparts were assembled using Network Brick approach^**21**^ and the whole circuit was engineered in two plasmids and inserted into *E.coli* DH5αZ1 cells (Fig. 1B). Detailed description of the plasmids can be found in supplementary table S1.

### Characterization of Feynman gate

Next, single colonies of the engineered DH5αZ1 were grown overnight in LB media with appropriate antibiotics, diluted and regrown in fresh LB media with combinations of inducers (aTc/IPTG) for 16 hours. The cells were spun down, washed and re-suspended in PBS for fluorescence measurements. Figure 2 shows that the expression pattern of the EGFP and E2Crimson in the ensemble (Fig. 2A) and in the single cell (Fig. 2B) as a function of aTc and IPTG and it matched appropriately with the Feynman gate truth table (Fig. 1A). We also built two other constructs (Fig S1A, B) for Feynman gate, where the relative copy numbers of various promoters and transcription factors were different. We observed that the experimental behaviors of those constructs were not optimum (Fig S2 A, B). We optimized the behavior by changing the relative copy numbers of various proteins and promoters by placing genetic cassettes in two plasmids with different copy numbers (Fig. 1B).

**Figure 2:**
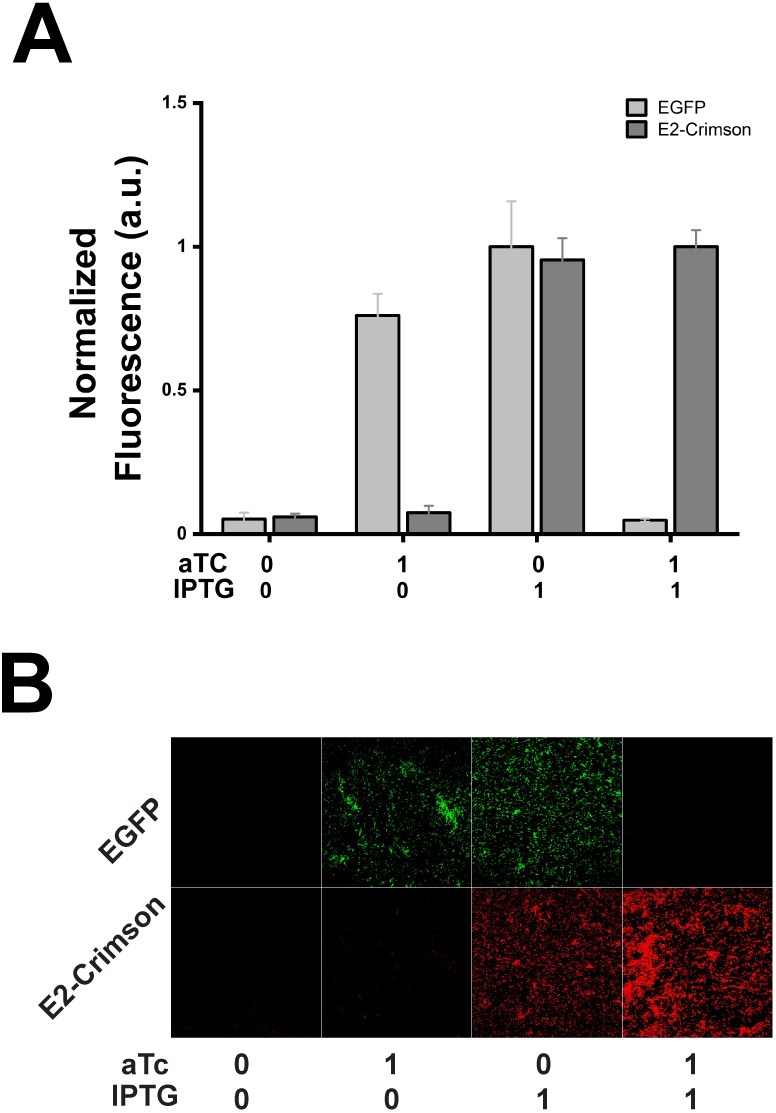
Engineered *E.coli* shows Feynman gate behavior: **A)** Ensemble level characterization of the device by measuring normalized EGFP and E2-crimson fluorescence against different induction states of input chemicals. Induction state 0 represents absence of the inducer molecules while 1 represents presence of inducers molecules at saturated concentrations ([aTc] = 200 ng/ml, [IPTG] = 10 mM). **B)** Single cell level characterization of Feynman gate behavior using fluorescence microscopy. The expressions of EGFP and E2Crimson were captured by separate ‘green’ and ‘red’ output channels (see methods for details).

### Transfer function model

Next, we developed a simple transfer function model for the Feynman gate to capture the essential features of the circuit. The rate of accumulation of the fluorescent protein outputs within bacterial cells can be written as:

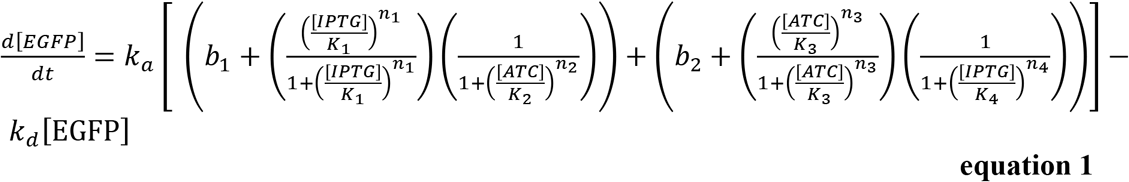

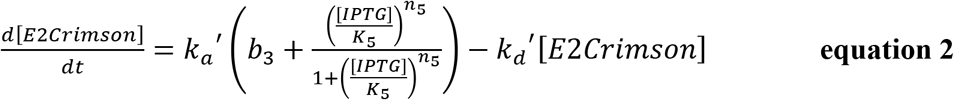

Where, *k_a_* and *k_a_′* are the scaling rate constants, which represents maximum additional production rate of EGFP and E2crimson respectively, due to up-regulation; *Ki* and *n_i_* represent the Hill constant (concentration of input signal at which 50% of maximum output signal is obtained) and Hill coefficients (measurements of ultrasensitivity) respectively; *k_d_* and *k_d_′* represent degradation rate of EGFP and E2Crimson. The degradation terms included protein decay and dilution due to cell division. The *bi* is the basal level expression from the respective promoters.

At steady-state, d[EGFP]/dT=0 and we can rearrange equations (1) as,

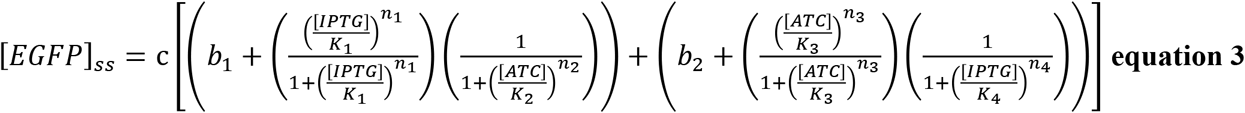

Where, 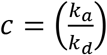

Similarly, at steady state d[E2Crimson]/dT=0, we can rearrange equations (2) as,

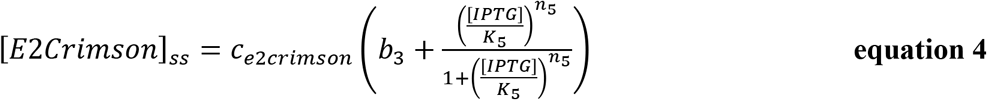

Where, 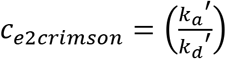

The equations and its parts represented specific genetic modules of the whole Feynman gate genetic circuit. The first and the second parts of the equation 3 represented two genetic modules for EGFP output module 1 (P_LAACCC_ –EGFP, P_TATA_ CIm, (in DH5alphaZ1) and module 2 (P_LCCTT_ EGFP, P_TATA_ CIm in DH5alphaZ1), respectively. Equation 4 represents E2Crimson output from P_Llaco-1_ –E2Crimson module.

### Dose response and Parameter Estimation

To estimate the parameter values in the transfer function model, we performed dose response experiments, where the output expression of EGFP and E2-Crimson were measured as a function of input IPTG and aTc concentration. We performed five of such dose response experiments (Fig. 3) to obtain all parameter values. In each dose response experiment, we varied the concentration of one inducer while keeping the other at the constant. The equation 3 was reduced to four equations at four different conditions.

**Figure 3:**
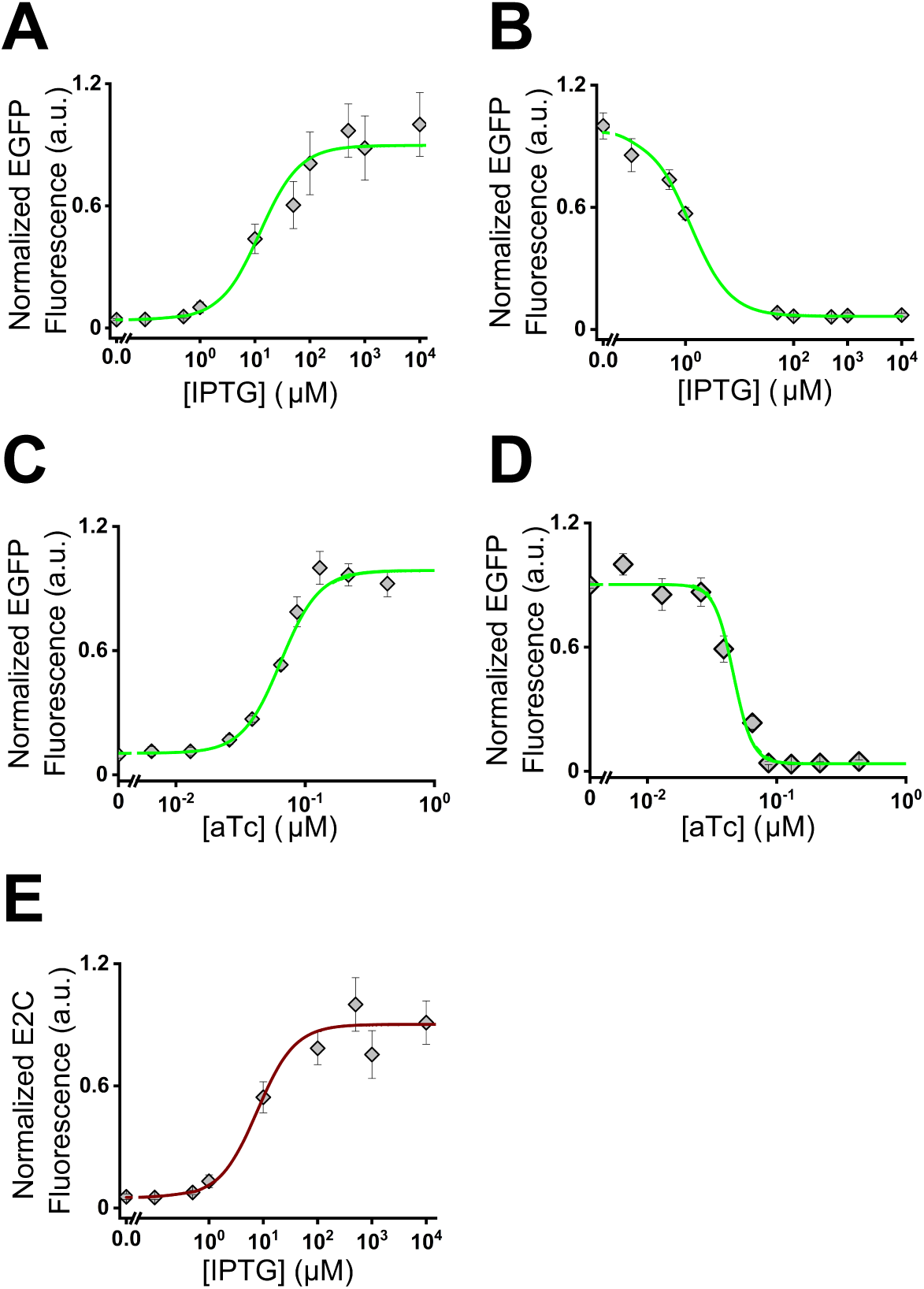
Dose response experiments of Feynman gate: Normalized fluorescence from EGFP and E2-crimson expression from the engineered *E.coli* were measured as a function of one inducer while keeping other inducer at its zero or saturated concentration. The data were fitted (solid line) using equations 5–8 and 4 (see main text). **A)** EGFP expression as a function IPTG with [aTc] = 0. **B)** EGFP expression as a function IPTG with [aTc] = 200 ng/ml. **C)** EGFP expression as a function aTc with [IPTG] = 0. **D)** EGFP expression as a function aTc with [IPTG] = 10mM. E) E2-Crimson expression as a function IPTG with [aTc] = 0.

When [aTc] = 0, we can rewrite eqn. (3) as:

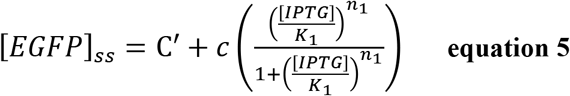

When [aTc] was kept at its saturated concentration (200ng/ml), in eqn. (3); 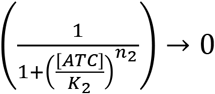 and this reduced the equation 3 into

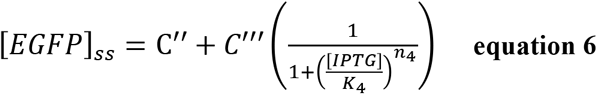

Similarly, When [IPTG] = 0, eqn. (3) becomes:

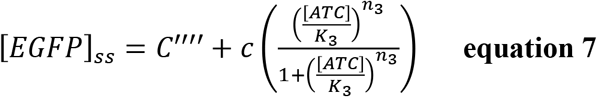

and at saturated concentration of IPTG,

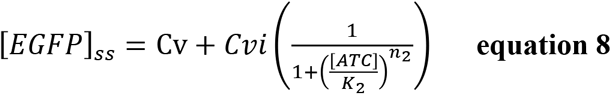

Where C(i)s are constants.

We measured EGFP fluorescence by performing dose response experiments (Fig. 3 A-D) at four specific inducer conditions as mentioned in equation 5–8.

IPTG dose response experiments were performed by keeping aTc concentration at zero (eq. 5) and at saturated concentration (eq. 6) and the aTc dose response experiments were performed by keeping IPTG concentration at zero (eq. 7) and at saturated concentration (eq. 8) (Fig. 3 A-D). Next, we fitted those experimental data with equations 5,6,7, and 8 as appropriate. Similarly, we measured E2crimson fluorescence as a function of IPTG concentrations while keeping aTc at zero concentrations and the dose response experiment was fitted with equation 4 (figure 3E). All the fitting values were tabulated in table S2. The Hill coefficients (ni) for all the dose response curves were found greater than 1. This (n>1) suggested that the synthetic genetic Feynman gate worked in an ultrasensitive or digital like way^**22**^. Next, by using all the parameter values in equation 3 and 4, we performed computational simulation for the genetic Feynman gate (Fig. 4A-C), which gave 3-dimensional topologies of the expression patterns of EGFP and E2-crimson as a function of IPTG and aTc. To experimentally validate the prediction from the simulation, we ran a set of experiments by simultaneously varying the concentration of ATC and IPTG and measured the EGFP and E2-Crimson fluorescence (Fig. 4D-F). The simulation and experiments showed a good topological match.

**Figure 4:**
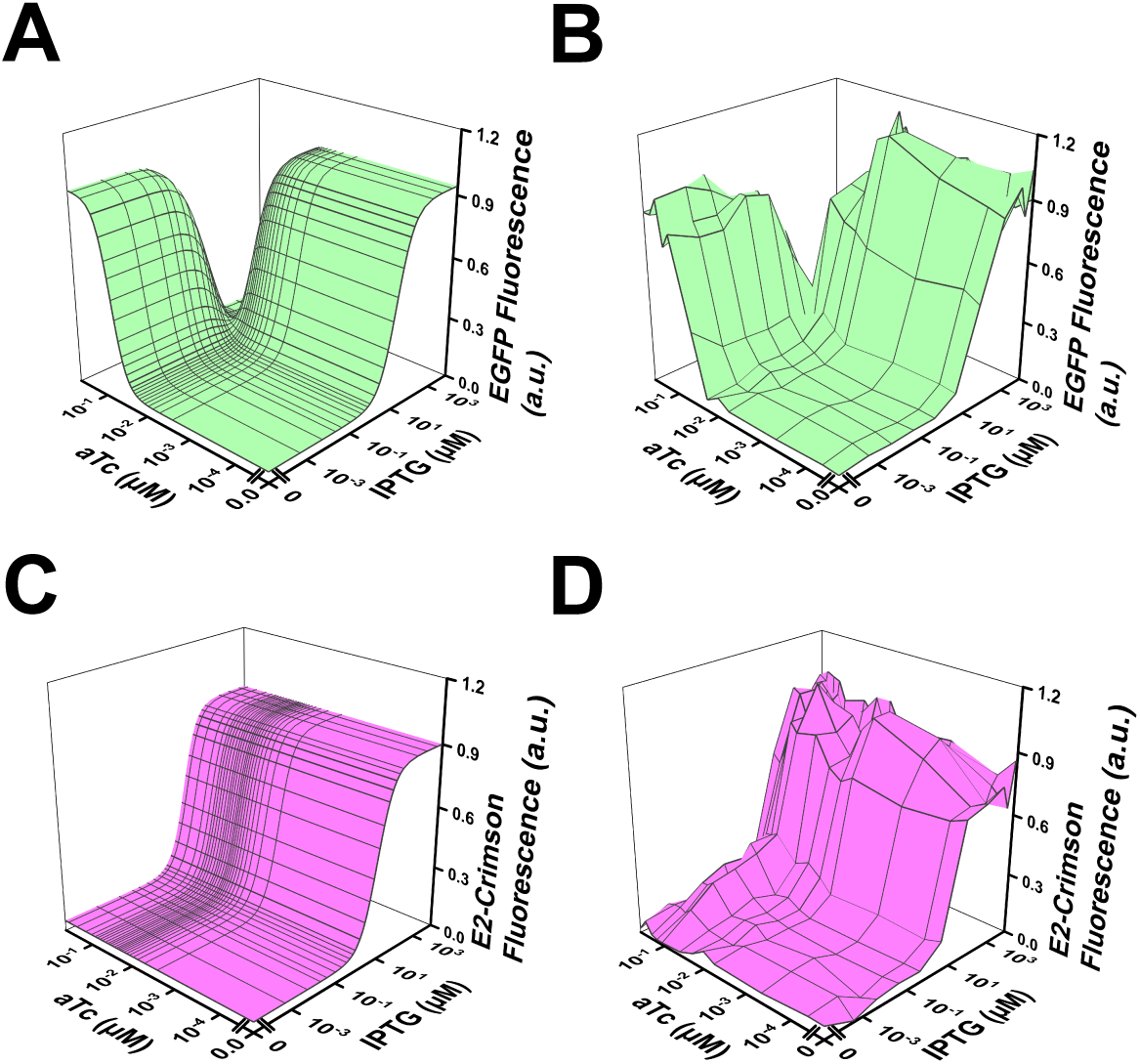
Simulation of genetic Feynman gate and its experimental validation. **A)** Simulated behavior of the device for the EGFP output **B)** Experimental behavior of the device for EGFP output. **C)** Simulated behavior of the device for the E2-crimson output. **D)** Experimental behavior of the device for E2-crimson output.

### Application of Feynman Gate in bacterial to mammalian cell information transfer

Next, we showed a simple application of the Feynman gate. Here we demonstrated shRNA-mediated gene silencing using Feynman gate by creating a bacterial cell-mammalian cell intercellular Feynman gate, where information was transferred from bacterial to the mammalian cell. The input signals (IPTG, aTc) were applied to the engineered invasive *E.coli* cells and the input information from bacteria was transferred through shRNAs to the mammalian (HeLa) cells, where the output signals were observed by the change in expression of two native genes CTNNB1 and AKT1. Here, we created the mammalian cell invasive *E.coli* by expressing invasin gene of *Yersinia enterocolitica*^**23**^ under a P_lux*_ promoter^4^ and hlyA genes of *Listeria monocytogenes*^**23**^ under a constitutive P_R_ promoter in a low copy plasmid (invasive module) in DH5alphaZ1 cells (Fig 5A). The invasin proteins expressed on the bacterial cell walls, which bind with beta 1-integrin expressing human cells and got engulfed^**23**^. The expressed hlyA gene inside the *E.coli* helped the engulfed *E.coli* to escape the endosomes^**23**^. In the invasion module, there was no luxR gene and its inducer AHL applied to the cell. In this situation the P_lux*_ promoter was not induced and allowed only a basal level transcription. We found that such un-induced basal level production of invasin was good enough to invade HeLa cell. Next, we re-engineered the synthetic genetic Feynman gate by replacing the two fluorescent output proteins with two shRNAs, anti-CTNNB1 shRNA and anti-AKT1 shRNA, which bind to human CTNNB1and AKT1 mRNAs, respectively and down regulate their expression in HeLa cells (Fig. 5A). Here we assembled all the bioparts for Feynman gate in a single plasmid. Both the invasion module and Feynman gate were incorporated in appropriate plasmids and inserted in *E.coli* DH5alphaZ1 cells. Before, reaching this final design, we performed a set of preliminary experiments to determine and optimize the behavior of the engineered *E.coli* and its interaction with the HeLa cell (Fig. S3).

**Figure 5:**
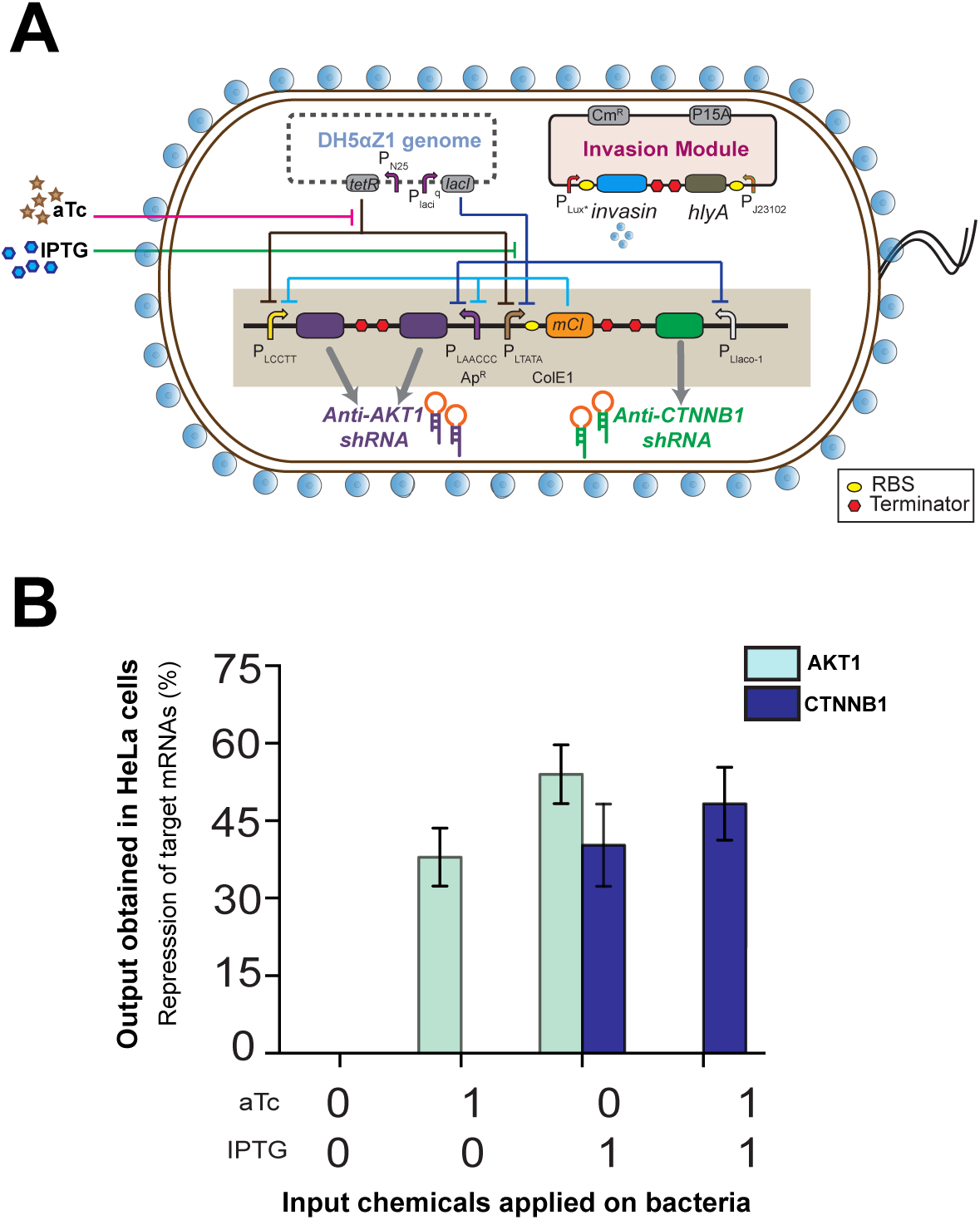
Bacterial-mammalian cell intercellular Feynman gate. **A)** Genetic circuit design of invasion module and Feynman gate in *E.coli*. **B)** Experimental behavior of the intercellular Feynman gate when environmental inputs (aTc and IPTG) applied on engineered bacteria and the outputs were manifested into endogenous gene silencing in HeLa cells. Induction state ‘0’ represents absence of the inducer molecules while ‘1’ represents presence of inducers molecules at saturated concentrations ([aTc] = 200 ng/ml, [IPTG] = 10 mM). The engineered invasive bacteria were co-cultured with HeLa cells for 2 hours. qRT-PCR analysis was performed after total mRNA isolation from HeLa cells harvested 24 hours post infection.

The engineered *E.coli* cells (Fig 5A) were grown with various combinations of aTc and IPTG, washed and co-cultured with HeLa cells (please see method section for details). As a control, we co-cultured HeLa with *E.coli* cells without any plasmids. After the co-culture, *E.coli cells* were washed off and mRNAs were extracted from the HeLa cells. Next, we measured the mRNA expression of AKT1 and CTNNB1 using RT-PCR. The result showed a matching Feynman gate behavior, where the input chemical logics were applied in the bacterial cells and the output logics were manifested within HeLa cell as relative repressions of the AKT1 and CTNNB1 mRNAs (Figure 5B).

Such information transfer of reversible logic between bacteria and human cells may have implications in programmed drug delivery. Given that the input and outputs were mapped in a one to one fashion in reversible computing, synthetic genetic reversible logic gates may have implications in sensing the patterns of multiple chemicals in a sample. Such systems may have applications in precision diagnostics by identifying the patterns and compositions in intracellular chemical states just by observing the outputs. Similarly, reversible logic gates may have applications as chemical sensors in environments and industry, where the complex pattern of presence and absence of multiple chemicals can be observed.

In conclusion, we created a 2-input-2-output synthetic genetic reversible Feynman logic gate in single *E.coli* cells. Reversible logic gates are the main components of reversible computing, where input and outputs were mapped in a unique one to one pattern, such that the pattern of input signal can be read from the output signals. We used extracellular IPTG and aTc as input signals and the expression of EGFP and E2-crimson gene as the output signals. The circuit was built and optimized in *E.coli* DH5αZ1 cells using network brick approach. A simple transfer function model was developed and simulation was performed to capture the basic features of the circuit. We demonstrated that the genetic Feynman gate is ultrasensitive or digital-like and its behavior was predictive in nature. We further showed an application of the Feynman gate, where input information from bacteria was computed and transferred to HeLa cells and the output signals were observed as expression changes of two native genes of HeLa cells. Given that one-to-one input-output mapping, such reversible genetic logic gates may have applications in diagnostics and sensing, where compositions of multiple intracellular or environmental chemicals could be estimated by observing the outputs.

## Methods

### Plasmid Construction

Plasmid vectors with NetworkBrick^**21**^ assembly module were used for plasmid construction. All plasmids are listed in table S1. The bioparts (promoter, ribosome binding sites, gene and transcription terminators) were arranged within appropriate network brick empty vectors using standard molecular cloning protocols. The individual expression cassettes were then assembled using network brick assembly process. A detailed process pipeline can be found elsewhere^**21**^. PCR amplifications were performed using KOD Hot Start DNA polymerase (Merck Millipore). All restriction enzymes, T4DNA ligase, and DNA ladders were obtained from New England BioLabs. Plasmid isolation, gel extraction, and PCR purification kits were obtained from QIAGEN. All primers and oligos were synthesized from Integrated DNA Technologies, Inc. Invasin and HlyA gene products were custom synthesized. Sequence of bioparts and oligos were shown in table S3. Sequencing of cloned genes and bio-parts in plasmid constructs were carried out by Eurofins Genomics India Pvt. Ltd.

#### Bacterial strains and Growth

Chemically competent *E.Coli*. DH5αZ1 cells were co-transformed with appropriate plasmids (Fig 1B). Distinct and isolated single colonies from LB-Agar (Difco,Beckton Dickinson) plates were picked for overnight cultures. Luria–Bertani (LB) media (HiMedia) with appropriate antibiotics (100 μg/ml and 34 μg/ml for ampicillin (HiMedia) and chloramphenicol (HiMedia) respectively) was used for bacterial growth. Cells for plasmid isolation were grown overnight at 37°C. For conducting experiments, subculture was done by 100-fold dilution in fresh LB media with antibiotics and inducers IPTG ( Sigma Aldrich) or aTc (200 ng/ml) at appropriate concentration. Cells were grown for 16 hr at 37°C with shaking at ~250 RPM, pelleted down and washed thrice before fluorescence measurements. For infection assays, cells were cultured in BHI broth (Himedia) at 37°C, with shaking at ~250rpm followed by 1:100 dilution and subculture in fresh BHI broth with appropriate antibiotics and inducers.

#### shRNA design and Construction

Each shRNA (anti-shAKT1^**24**^, anti-shCTNNB1^**23**^, and other shRNAs, table S3) was designed so that the sense strand of the shRNA stem was homologous to the target and the anti-sense strand of the shRNA stem was complementary to the target. Sense and antisense strands were connected by a 9-nt loop and two ‘T’ residues were added in the 3’-end. Forward and reverse complementary strands of the hairpin oligos were flanked by EcoRI and XbaI at respective 5’ and 3’ends and were custom synthesized. Sequences of the shRNAs were shown in table S3. Each shRNA insert was constructed by annealing complementary oligos using standard protocol to create a synthetic DNA insert that was cloned into network brick plasmids under appropriate promoters. The shRNA expression cassettes consisted of the promoter, hairpin and terminator.

#### Fluorescence and optical density (OD) measurements

Cells from 16 hours cultures with various concentrations of inducers as appropriate were washed thrice and resuspended in PBS before taking the spectroscopic measurements. All experimental data were collected from minimum of 3 independent colonies. DH5αZ1 cells carrying no plasmids were taken as the control. For the validation experiments, 10X10 or more intermediate concentration points for each of the two inducers were used. Cells were loaded onto 96-well plate (black, Greiner Bio-One) for fluorescence measurement using Synergy HTX Multi-Mode reader (Biotek Instruments, USA). For fluorescence measurements for EGFP, the cells were excited by a white light source that had been passed through an excitation filter 485/20 nm and emission was collected by 516/20 nm bandpass filter with appropriate gain. E2-Crimson fluorescence was measured using an excitation 610/10 nm bandpass filter and emission 645/10 nm bandpass filter. The OD_600_ was also measured in the same instrument.

#### Data Analysis, Curve Fitting and Simulation

Fluorescence normalization was performed by dividing the raw fluorescence values by respective OD_600_ values. Auto-fluorescence was measured as average normalized fluorescence of the DH5αZ1 cells (no plasmid) and subtracted from the normalized fluorescence value of the experimental set. This gave the unscaled normalized fluorescence values. Data were taken for at least 3 biological replicates for each condition.

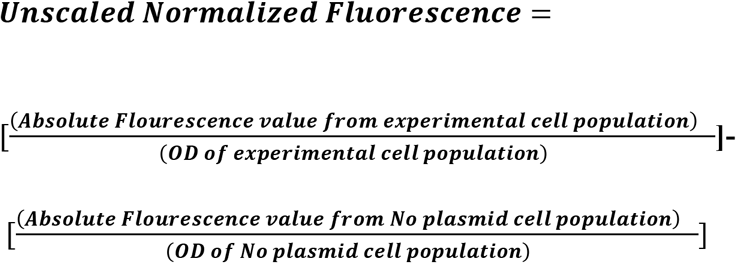

Unscaled values thus obtained were scaled down between 0 and 1, considering the normalized fluorescence value at induction point of maximum expected fluorescence to be ‘1’ using the following formula.

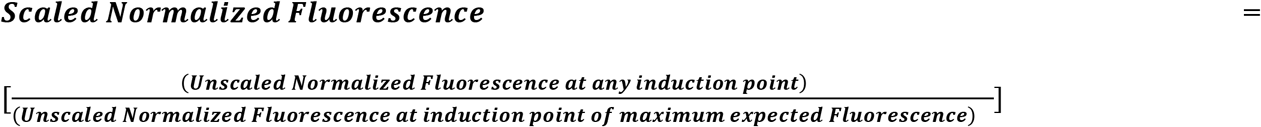

Scaled fluorescence data from Fluorescence and optical density (OD) measurements were plotted and analyzed using OriginPro 2018 (OriginLab Corporation, USA). The dose response curves were fitted with the appropriate Hill equations as discussed in results and discussion using built-in Levenberg Marquardt algorithm. The simulations were performed by using parameterized equations and by generating calculated fluorescence values against simultaneously varying concentrations of two inducers across 50X50 points.

#### Confocal Microscopy

Bacterial cells were pelleted down out at 4500RPM for 3 minutes from the 16 hours subcultures and washed trice in PBS. Washed cells were resuspended in PBS. To visualize, 10μl cell suspension was placed over 1% percent level agarose pad (SeaKem LE agarose). The samples were then observed under 60X water immersion objective (NA: 1.4) in Nikon AIRsi confocal microscope equipped with the resonant scanner and coherent CUBE diode laser system. The samples were subjected to excitation of the fluorescent proteins by 488 nm and 561 nm Lasers channels and their emissions (emission filters BP 525/50 nm and BP 585/65 nm) were captured using NIS Elements Imaging Software. Raw microscopic images were processed using Fiji software.

#### Bacterial and mammalian cell co-culture and infection

Chemically competent *E.Coli*. DH5αZ1 cells were co-transformed with shRNAs bearing plasmid and invasion module plasmid. This served as our experimental infection strain. We used *E.Coli*. DH5αZ1 cells without any plasmid as control cells for the infectivity experiments. Isolated single colonies from LB agar plates were picked and overnight culture was given in Brain-Heart-Infusion broth with ampicillin and chloramphenicol for infection strain and without antibiotics for control *E.coli*. The engineered cells were then subcultured after 1:100 dilutions in fresh BHI broth with ampicillin and chloramphenicol (or no antibiotics for control *E.coli*) and appropriate inducers for 10 hours. Before infecting HeLa cells, bacterial cells were pelleted down at 4500 RPM at 4°C for 15 min, washed twice, resuspended in DMEM medium without serum and with ampicillin and chloramphenicol (or no antibiotics for control *E.coli*) and grown for another 2 hours with appropriate inducers. We defined this as the infection media. HeLa cells were cultured in DMEM medium (GIBCO) with 10% FBS (GIBCO) supplemented with 100 U/ml penicillin G, 10 mg/ml streptomycin and 2.5 mg/ml amphotericin (Sigma). For bacterial invasion, HeLa cells were cultured in the 60×15mm dishes (Nunc) at around 30% confluency two days before adding engineered bacteria. 30 minutes in prior to bacterial infection, the HeLa cell culture medium was replaced with fresh serum-free DMEM medium with 100 U/ml penicillin G, 10 mg/ml streptomycin and 2.5 mg/ml amphotericin. Early log phase engineered bacterial cells in infection media were then added to the serum starved HeLa cells at the 1000:1 MOI (the ratio of infectious agents (here engineered *E.coli*) to infection targets (here HeLa cell)). MOI was calculated based on approximate number of bacterial cells from OD_600_ values and approximate confluency of Hela cells. After two hours of infection exposure, HeLa cells were washed extensively and treated with 150 mg/ml of gentamycin and other antibiotics in DMEM media with FBS for 30 minutes. The HeLa cells were then washed thrice and cultured in DMEM media with FBS and appropriate antibiotics for additional 24-h in a CO_2_ incubator (Eppendorf Galaxy 170R) before harvesting for downstream processing.

#### RT-PCR

Total RNA was isolated from infected Hela cells (experiment) and HeLa cells co-cultured with non-engineered *E.coli* (control) using RNeasy Mini Kit (Qiagen) and then reverse transcribed using QuantiTect Reverse Transcription Kit (Qiagen) following the manufacturer’s protocol. All oligos used for qRT-PCR are listed in table S3. Then qRT-PCR was performed using QuantiTect-SYBR GREEN PCR Kit (Qiagen) in StepOne Real-Time PCR system (Applied Biosystems) with B-actin as housekeeping standard. At least 3 biological replicates and three technical replicates were used for the study. The relative mRNA expression levels were measured using standard 2^−ΔΔCt^ method.

## Acknowledgements

This work was financially supported by the grant RSI4002 (Department of Atomic Energy, Govt. of India) and SERB (CRG/20lB/00l394), Govt. of India

## Author Contributions

SB conceived and designed the study. RS performed all of the experiments. KS and DB performed some initial experiments. RS and SB designed the experiments, analyzed and interpreted the data, and wrote the paper.

