## Supplementary Figures and Tables for "A Synthetic Genetic Reversible Feynman Gate in a Single *E.coli* Cell and its Application in Bacterial to Mammalian Cell Information Transfer"

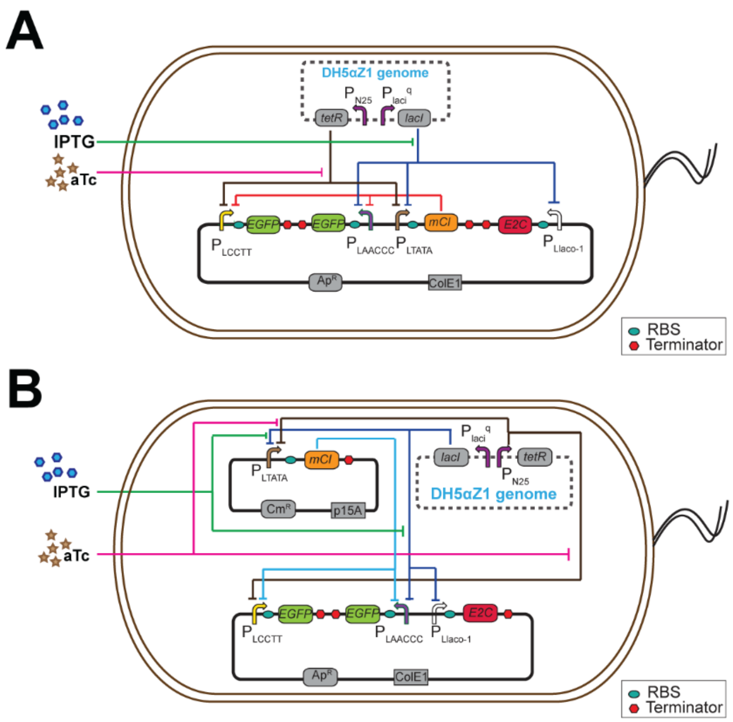


**Figure S1: Design-2 and Design-3 for biological 2x2 Feynman Gate in engineered *E. coli*.** A) Diagram depicting Design-2 for the biological Feynman Gate. Here, all the gene expression cassettes for the reporter and regulatory proteins are assembled over a single medium copy (ColE1) plasmid in alternate direction. B) Diagram depicting Design-3 for the biological Feynman Gate. Here, all the expression cassettes for the reporter protein are assembled over a medium copy (ColE1) plasmid in alternate direction. The cassette expressing mCI gene is placed in a low copy (P15A) plasmid.

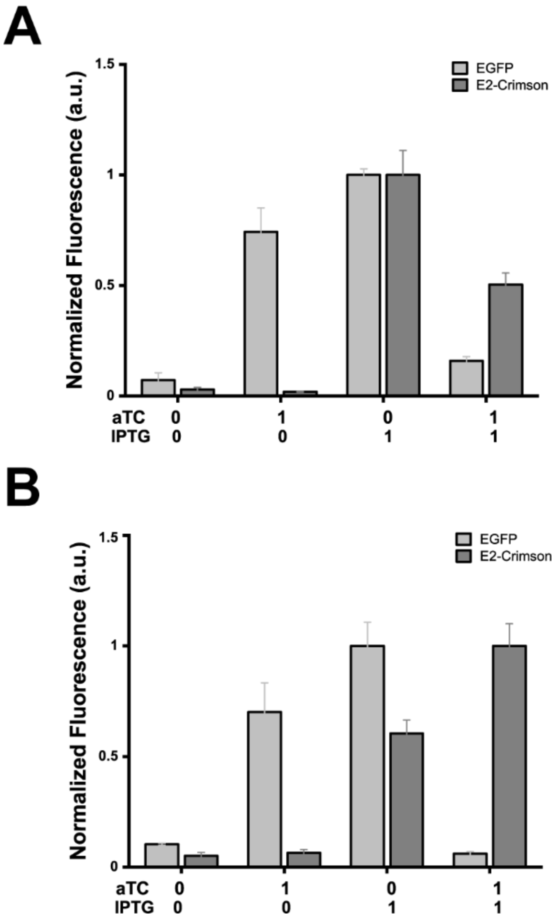


**Figure S2: Characterization of Design-2 and Design-3 for biological Feynman Gate** A) Characterization of Design-2 fabricated in *E.coli* DH5αZ1 by measuring normalized EGFP and E2-crimson fluorescence against different induction states of input chemicals. Induction state 0 represents absence of the inducer molecules while 1 represents presence of inducers molecules at saturated concentrations ([aTc] = 200 ng/ml, [IPTG] = 10 mM). B) Characterization of Design-3 fabricated in *E.coli* DH5αZ1 by measuring normalized EGFP and E2-crimson fluorescence against different induction states of input chemicals. Induction state 0 represents absence of the inducer molecules while 1 represents presence of inducers molecules at saturated concentrations ([aTc] = 200 ng/ml, [IPTG] = 10 mM).

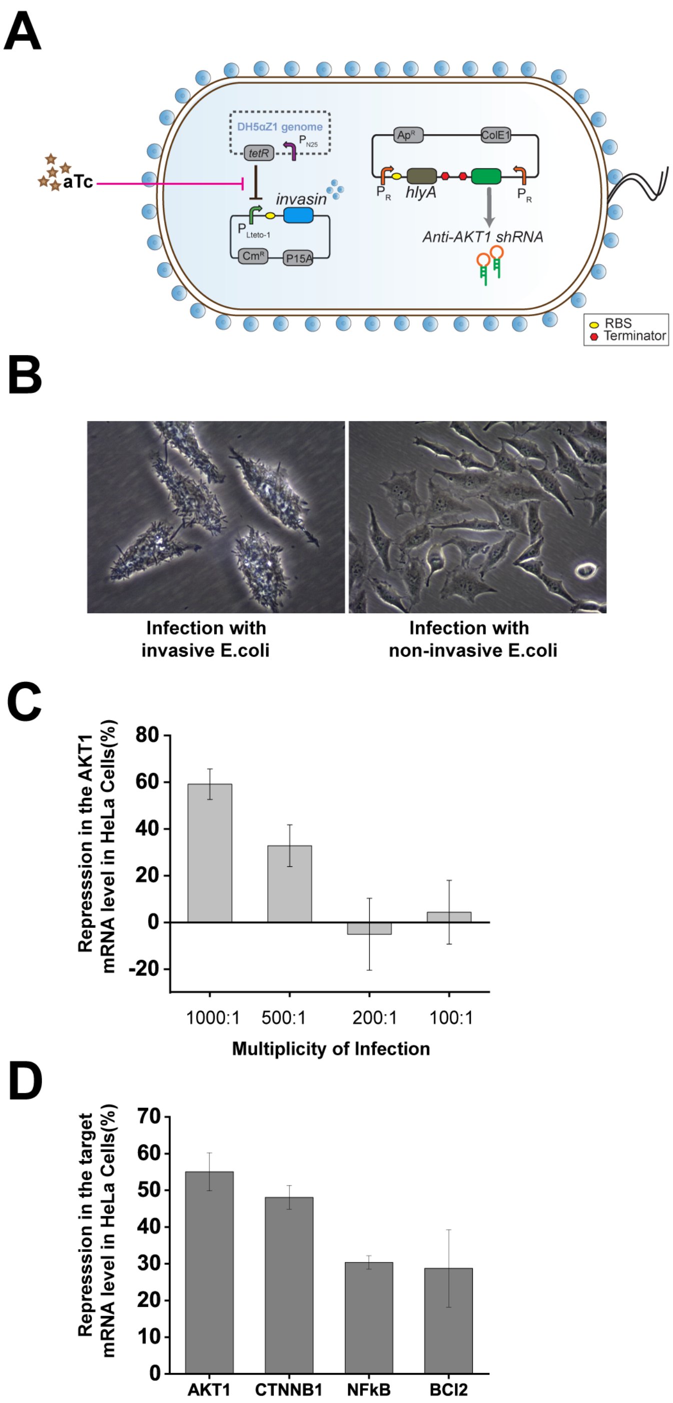


**Figure S3: Engineered bacteria invades HeLa cells and silences endogenous target gene** A) Diagram depicting the design of invasion and delivery module integration with Anti-AKT1 shRNA in *E. coli*. The invasin gene is placed under a aTc inducible promoter while hlyA and Anti-AKT1 shRNA is placed under a constitutive Pr promoter. The device receives aTc ([aTc] = 200 ng/ml) as chemical inducer synthesizes invasin which mediates bacterial internalization to b1-integrin expressing HeLa cells expresses. B) Bright field image of HeLa cells infected with engineered invasive *E.coli* (Left) and non-engineered non-invasive *E.coli* cells (Right). C) Effect of MOI on repression in the mRNA level of AKT1 gene in HeLa cells. D) Repression in the mRNA level of respective endogenous target genes in HeLa cells upon infection with invasive bacteria loaded with Anti-AKT1, Anti-CTNNB1, Anti-NFkB, and Anti-BCl2 shRNAs. HeLa cells were incubated for 24 hours after infection with engineered bacteria at MOI 1000:1.

**Table S1: Description of Plasmids**

| Sr no | Plasmid Name | Origin of replication | Antibiotic selection | Description | Source |
|---|---|---|---|---|---|
| **1** | pOR-*EGFP*-12 | ColE1 | Amp | Source of *EGFP*, Ampicillin resistance, ColE1 origin | Kind gift from Prof. David McMillen, UofT |
| **2** | pOR-Luc-31 | P15A | Cmr | Source of $P_L$tetO-1 and Chloramphenicol resistance, P15A origin | Kind gift from Prof. David McMillen, UofT |
| **3** | pNB_A_FCEV | p15A | Amp | Empty network brick vector for assembly in forward direction | [Literature Ref 21] |
| **4** | pNB_E_FCEV | ColE1 | Amp | Empty network brick vector for assembly in forward direction | |
| **5** | pNB_U_FCEV | pUC | Amp | Empty network brick vector for assembly in forward direction | |
| **6** | pNB_RCEV | ColE1 | Amp | Empty network brick vector for assembly in Reverse direction | |
| **7** | pPINA7A2EGFP(F) | ColE1 | Amp | Source of PLAACCC promoter | [Literature Ref 18] |
| **8** | pPANI2A2EGFP | ColE1 | Amp | Source of CCTT promoter | |
| **9** | pPIAA7A1EGFP | pUC | Amp | Source of TATA promoter | |
| **10** | pRA1SEGFPTcIfm | pUC | Amp | Source of frame-shifted CI gene | [Literature Ref 20] |
| **11** | pNB_EA_$P_{lcctt}$EGFP_F | ColE1 | Amp | EGFP expression cassette (Forward) | This study |
| **12** | pNB_EA_$P_{laaccc}$EGFP_R | ColE1 | Amp | EGFP expression cassette (Reverse) | This study |
| **13** | pNB_EA_$P_{ltata}$mCI_F | ColE1 | Amp | mCI expression cassette (Forward direction) | [Literature Ref 4] |
| **14** | pNB_EA_$_{Pllaco-1}$E2crimson | ColE1 | Amp | E2-crimson expression cassette (Reverse) | This study |
| **15** | pNB_AR_$_{Pllaco-1}$e2crimson | P15A | Cmr | Plasmid-1. E2-Crimson expression cassette in design-1 under low copy plasmid (Forward) | This study |
| **16** | pNB_EA_$P_R$INV | ColE1 | Amp | Invasin under constitutive Pr promoter | This study |
| **17** | pNB_EA_$P_{lteto-1}$Inv | ColE1 | Amp | Invasin under inducible Plteto-1 promoter in medium copy | This study |
| **18** | pAR31inv | p15a | Cmr | Invasin under inducible Plteto-1 promoter in low copy | This study |
| **19** | pNB_EA_$P_R$HLYA | ColE1 | Amp | hlyA under constitutive Pr promoter | This study |
| **20** | pNB_AR1INVPrHly | p15A | Cmr | Pooled Plteto-1 invasin and Pr hlyA in forward and reverse direction | This study |

| | | | | | |
|---|---|---|---|---|---|
| **21** | pNB_EA_$_{Pllaco\text{-}1}$CTNNB1 | ColE1 | Amp | Anti-CTNNB1 shRNA under a inducible Pllaco-1 promoter in medium copy | This study |
| **22** | pNB_AR_$P_{llaco\text{-}1}$CTNNB1 | P15A | Cmr | Anti-CTNNB1 shRNA under a inducible Pllaco-1 promoter in low copy | This study |
| **23** | pNB_EA_$P_{lcctt}$AKT1 | ColE1 | Amp | Anti-AKT1 shRNA under a inducible Plcctt promoter | This study |
| **24** | pNB_EA_$P_{laaccc}$AKT1 | ColE1 | Amp | Anti-AKT1 shRNA under a inducible Plaaccc promoter | This study |
| **25** | pNB_EA_$P_{J23}$HLYA | ColE1 | Amp | hlyA under constitutive $P_{J23102}$ promoter | This study |
| **26** | pNB_EA_$P_R$AKT1 | ColE1 | Amp | Anti-AKT1 shRNA under constitutive promoter | This study |
| **27** | pNB_EA_$P_R$CTNNB1 | ColE1 | Amp | Anti-CTNNB1 shRNA under constitutive promoter | This study |
| **28** | pNB_EA_$P_{lcctt}$EGFP_F $P_{laaccc}$EGFP_R | ColE1 | Amp | Pooled PlccttEGFP and PlaacccEGFP expression cassettes in forward and reverse direction repectively | This study |
| **29** | pNB_EA_$P_R$HlyA_F_$P_R$CTNNB1 | ColE1 | Amp | Pooled PrHLYA and PrShCTNNB1 cassettes in forward and reverse direction respectively | This study |
| **30** | pNB_EA_$P_{lcctt}$EGFP_F_$Pl_{aaccc}$EGFP_$Pl_{tata}$mCI_F | ColE1 | Amp | Plasmid-2. Pooled PlccttEGFP, PlaacccEGFP , and PltataCI cassettes in forward, reverse and forward direction respectively | This study |
| **31** | pNB_EA_PlccttEGFP_F_$Pl_{aaccc}$EGFP_$Pl_{tata}$mCI_F_$Pl_{laco\text{-}1}$E2crimson_R | ColE1 | Amp | Pooled PlccttEGFP, PlaacccEGFP , and PltataCI, Pllaco-1-E2crimson, cassettes in forward, reverse, forward and reverse direction respectively | This study |
| **32** | pNB_EA_PlccttEGFP_F_$Pl_{aaccc}$EGFP_$Pl_{laco\text{-}1}$E2crimson_F | ColE1 | Amp | Pooled PlccttEGFP, PlaacccEGFP, Pllaco-1E2crimson caessettes in forward, reverse and forward direction respectively | This study |
| **33** | pNB_EA_$Pl_{cctt}$AKT1_F_$Pl_{aaccc}$AKT1 | ColE1 | Amp | Pooled PlccttShAKT1, PlaacccShAKT1 in forward and reverse direction respectively | This study |
| **34** | pNB_EA_PlccttAKT1_F_$Pl_{aaccc}$AKT1_R$Pl_{tata}$mCI_F | ColE1 | Amp | Pooled PlccttShAKT1, PlaacccShAKT1 and PltatamCI cassettes in forward, reverse and forward direction respectively | This study |
| **35** | pNB_EA_$Pl_{cctt}$AKT1_F_$Pl_{aaccc}$AKT1_R$Pl_{tata}$mCI_$Pl_{laco\text{-}1}$CTNNB1_R | ColE1 | Amp | Pooled PlccttShAKT1, PlaacccShAKT1 PltatamCI, Pllaco-1ShCTNNB1 in forward, reverse, forward and reverse direction respectively | This study |
| **36** | pNB_EA_$P_R$NFkB | ColE1 | Amp | Anti-NFkB shRNA under constitutive promoter | This study |
| **37** | pNB_EA_$P_R$Bcl2 | ColE1 | Amp | Anti-Bcl2 shRNA under constitutive promoter | This study |

**Table S2: Curve fitting parameter values**: Dose response curves of Biological Feynman Gate for aTc and IPTG chemical signals yielded normalized fluorescence data which were fitted with appropriate equations.

| Parameter | Estimated value | Standard Error | Unit |
|---|---|---|---|
| **For EGFP expression** | | | |
| $b_1$ | 0.03 | - | - |
| $b_2$ | 0.03 | - | - |
| $K_1$ | 12.28677 | 2.26445 | μM |
| $K_2$ | 0.04619 | 0.00424 | μM |
| $K_3$ | 0.06464 | 0.00225 | μM |
| $K_4$ | 1.21834 | 0.14945 | μM |
| $n_1$ | 1.2 | 0.00 | - |
| $n_2$ | 5.9 | 1.38479 | - |
| $n_3$ | 2.9 | 0.25291 | - |
| $n_4$ | 1.17 | 0.17107 | - |
| c | 0.89 | - | - |
| **For E2-Crimson expression** | | | |
| $b_3$ | 0.06164 | 0.01119 | - |
| $K_5$ | 7.7571 | 2.20668 | μM |
| $n_5$ | 1.25133 | 0.21616 | - |
| $c_{e2crimson}$ | 0.85 | - | - |

**Table S3: List of promoters, primers, oligos and RBSs.**

| Primer name | Sequence (5' - 3') | Purpose |
|---|---|---|
| E2C_F | TGGATAGCACTGAGAACGTCAT | Amplification of fluorescent protein E2- Crimson: forward primer (1st round) |
| E2C_R | ACCACGGTGTAGTCCTCGTT | Amplification of fluorescent protein E2- Crimson: reverse primer (1st round) |
| PR_F | TACTGGGACGAAGACGAACA | Amplification of cloned cassette in network Brick plasmid: Forward primer |
| Sfx_R | GTTGTTTTGGAGCACGGAAC | Amplification of cloned cassette in network Brick plasmid: Reverse primer |
| Inv_F | GTTTGACGTATGACAGGTATGC | To amplify the Invasin gene: Forward primer,1$^{st}$ round |

| | | |
|---|---|---|
| Inv_R | TTATATTGACAGCGCACAGA | To amplify the Invasin gene: Reverse primer,1st round |
| Inv_exa_F | AGGGTAGAATTC GTTTGACGTATGACAGGTATGC | To amplify the Invasin gene: Forward primer, 2nd round |
| Inv_exa_R | AGGGTCTCTAGA AGGGTAGAATTC GTTTGACGTATGACAGGTATGC | To amplify the Invasin gene: Reverse primer, 2nd round |
| hlyA_F | ccctcctttgattagtatattcctatctta | To amplify the hlyA gene: Forward primer, 2nd round |
| hlyA_rp | aagcttttaaatcagcaggggtctttttgg | To amplify the hlyA gene: Reverse primer, 2nd round |
| AKT1_F | aattcTGCCCTTCTACAACCAGGAttcaagagaTCCTGGTTGTAGAAGGGCAtt | Anti-AKT1 oligo sequence (forward) |
| AKT1_R | **ctag**aaTGCCCTTCTACAACCAGGAtctcttgaaTCCTGGTTGTAGAAGGGCAg | Anti-AKT1 oligo sequence (Reverse) |
| CTNNB1_F | aattcAGCTGATATTGATGGACAGttcaagagaCTGTCCATCAATATCAGCTtt | Anti-CTNNB1 oligo sequence (forward) |
| CTNNB1_R | **ctag**AAAGCTGATATTGATGGACAGtctcttgaaCTGTCCATCAATATCAGCTg | Anti-CTNNB1 oligo sequence (reverse) |
| Egfp _F | CAAGGGCGAGGAGCTGTT | 1st round amplification of *EGFP* (Forward primer) and Forward Sequencing Primer for EGFP |
| Egfp_R | CCATGCCGAGAGTGATCC | 1st round amplification of *EGFP* (Reverse primer) |
| Bcl2_F | aattcGCTGCACCTGACGCCCTTCttcaagagaGAAGGGCGTCAGGTGCAGCtt | Anti-Bcl2 oligo sequence (forward) |
| Bcl2_R | ctagaaGCTGCACCTGACGCCCTTCtctcttgaaGAAGGGCGTCAGGTGCAGCg | Anti-Bcl21 oligo sequence (reverse) |
| NFkB_F | aattcCGCCCTATCCCTTTACGTCAttcaagagaTGACGTAAAGGGATAGGGCtt | Anti-NFkB oligo sequence (forward) |
| NFkB_R | ctagaaGCCCTATCCCTTTACGTCAtctcttgaaTGACGTAAAGGGATAGCGCGg | Anti-NFkB oligo sequence (reverse) |
| RBS | ATTAAAGAGGAGAAA | Sequence of RBS present in all expression cassettes except invasin |
| $RBS_i$ | tttcatttaaattatgatggtt | Sequence of RBS present in all invasin expression cassettes |
| qAKT1_L | CTTCTATGGCGCTGAGATTGT | Primer of amplification of AKT1 gene (Forward) |
| qAKT1_R | GCC CGA AGT CTG TGA TCT TAA T | Primer of amplification of AKT1 gene (Forward) |
| qBCL2_L | GTG GAT GAC TGA GTA CCT GAA C | Primer of amplification of AKT1 |

| | | |
|---|---|---|
| | | gene (Forward) |
| qBCL2_R | GAG ACA GCC AGG AGA AAT CAA | Primer of amplification of AKT1 gene (Forward) |
| qNFkB_L | CTG TCC TTT CTC ATC CCA TCT T | Primer of amplification of AKT1 gene (Forward) |
| qNFkB_R | TCC TCT TTC TGC ACC TTG TC | Primer of amplification of AKT1 gene (Forward) |
| qBACT_L | GGA CCT GAC TGA CTA CCT CAT | Primer of amplification of AKT1 gene (Forward) |
| qBACT_R | CGT AGC ACA GCT TCT CCT TAA T | Primer of amplification of AKT1 gene (Forward) |
| qCTNB1_F | CCTTTGTCCCGCAAATCATG | Primer of amplification of AKT1 gene (Forward) |
| qCTNB1_F | CGTACGGCGCTGGGTATC | Primer of amplification of AKT1 gene (Forward) |
| shAKT1 | UGCCCUUCUACAACCAGGAUUCAAGAGAUC CUGGUUGUAGAAGGGCAUU | Short-hairpin RNA sequence against AKT1 gene |
| shCTNNB1 | AGCUGAUAUUGAUGGACAGUUCAAGAGACU GUCCAUCAAUAUCAGCUUU | Short-hairpin RNA sequence against CTNNB1 gene |
| shNFkB | CGCCCUAUCCCUUUACGUCAUUCAAGAGAU GACGUAAAGGGAUAGGGCUU | Short-hairpin RNA sequence against NFkB gene |
| shBCL2 | GCUGCACCUGACGCCCUUCUUCAAGAGAGA AGGGCGUCAGGUGCAGCUU | Short-hairpin RNA sequence against BCL2 gene |
| $P_{lteto-1}$ | CTCGAGTCCCTATCAGTGATAGAGATTGAC ATCCCTATCAGTGATAGAGATACTGAGCAC ATCAGCAGGACGCACTGACCGAATTC | |
| $P_{llac0-1}$ | CTCGAGAATTGTGAGCGGATAACAATTGAC ATTGTGAGCGGATAACAAGATACTGAGCAC ATCAGCAGGACGCACTGACCGAATTC | |
| $P_{lcctt}$ | CCTCGAGTACCTCTGGCGGTGATAGATTACC TCTGGCGGTGATATTGACATCCCTATCAGT GATAGAGATACTGAGCACATCCCTATCAGT GATAGAGAGAATTC | |
| $P_{laaccc}$ | CTCGAGAATTGTGAGCGGATAACAATTGAC ATTGTGAGCGGATAACAAGATACTGAGCAC ATCTACCTCTGGCGGTGATAGATGATTACC TCTGGCGGTGATAGATGATTACCTCTGGCG GTGATAGAATTC | |
| $P_{ltata}$ | CTCGAGTCCCTATCAGTGATAGAGATTGAC ATTGTGAGCGGATAACAAGATACTGAGCAC ATCCCTATCAGTGATAGAGAGATAATTGTG AGCGGATAACAATTGAATTC | |
| $P_R$ | CTCGAGTAACACCGTGCGTGTTGACTATTTT ACCTCTGGCGGTGATAATGGTTGCATGTAC GAATTC | |
| $P_{lux*}$ | CTCGAGACCTGTAGGATCGTACAGGTTTAC GCAAGAAAATGGTTTGTTACTTTCGAATAA AGAATTC | |
| $P_{J23102}$ | CTCGAGTTGACAGCTAGCTCAGTCCTAGGT ACTGTGCTAGCGAATTC | |
